## Supplementary Tables and Figures for "Mechanism of threonine ADP-ribosylation of F-actin by a Tc toxin"

### Supplementary data

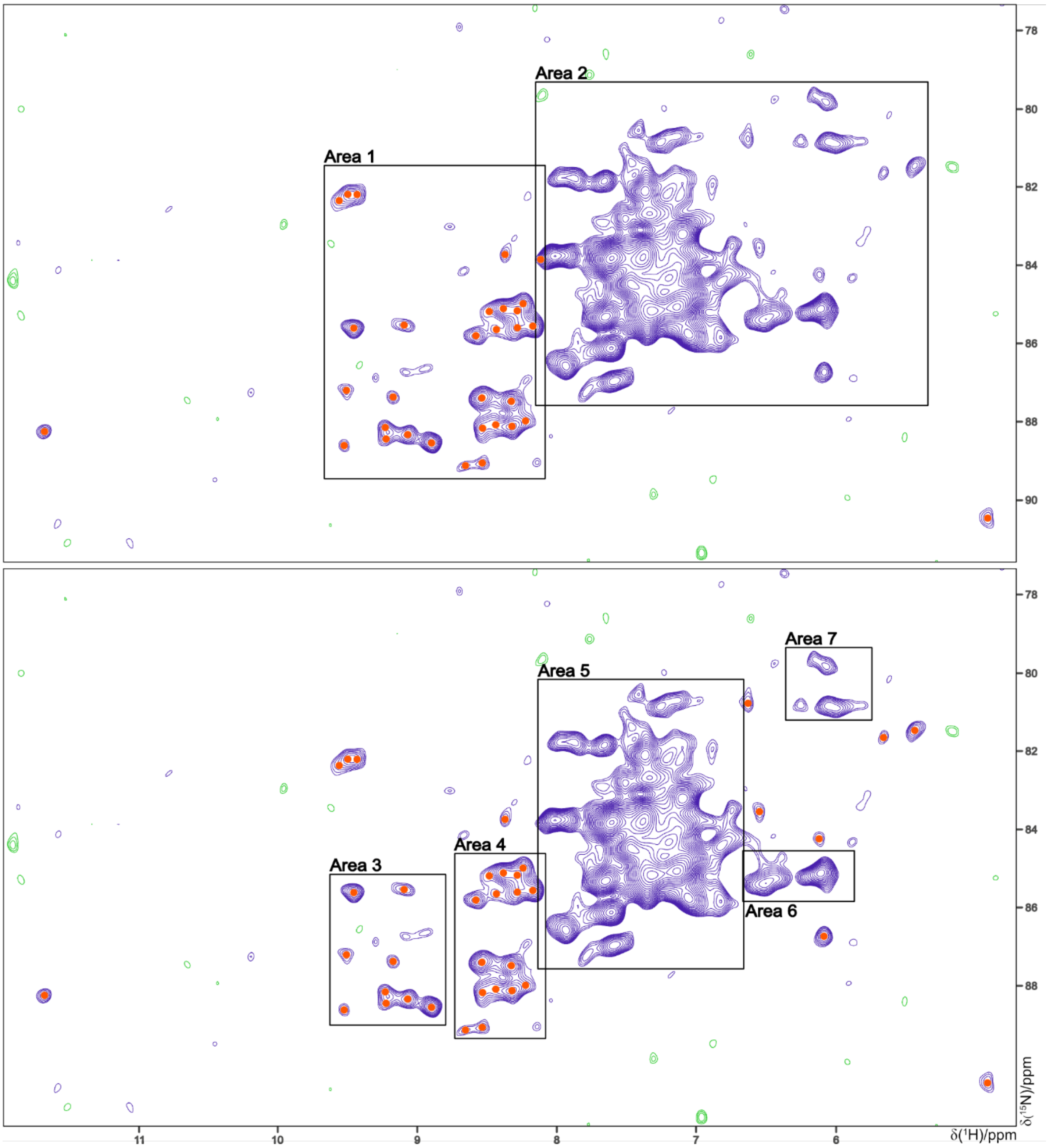

**Fig S1. Arginine NεH cross peak region of a two-dimensional $^{1}H$-$^{15}N$ correlation spectrum of the TcB-TcC cocoon** with two different signal integration schemes indicated by the frames. The different results obtained by counting the signals in Areas 1, 3, and 4 and taking the respective integrals as reference are listed in Table S1. The red dots indicate the counted peaks. In all other cases only the integrals of the areas were used in the calculations. The peaks outside the frames in both panels are counted individually.

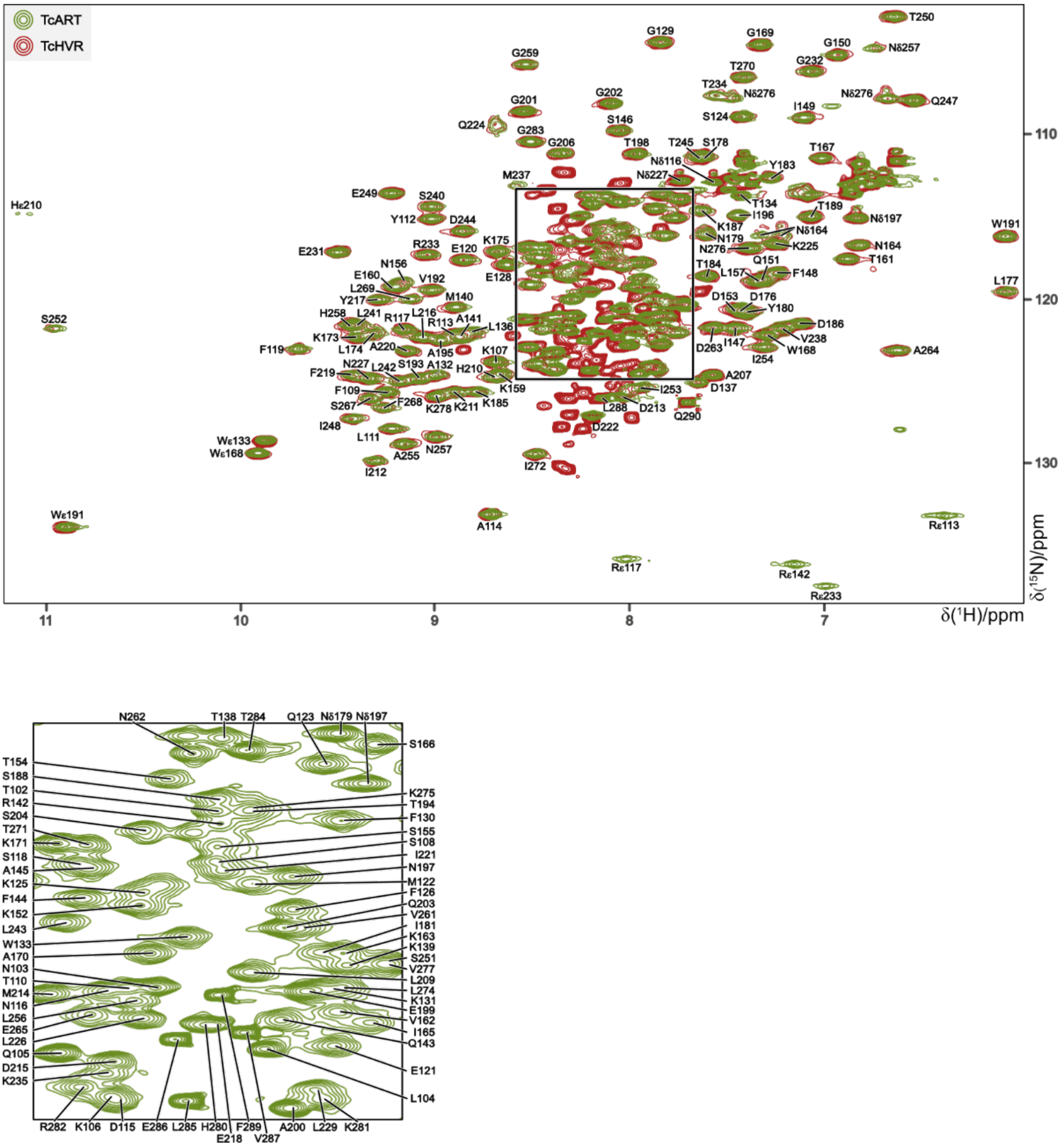

**Fig S2. Overlay of two-dimensional $^{1}$H-$^{15}$N correlation spectra of the TcHVR (red) and TcART (green) with assignments.** The framed region in the TcART spectrum on top (δ($^{1}$H): 7.7-8.6 ppm / δ($^{15}$N): 113.2-124.8 ppm) is shown below in an enlarged manner. The non-overlapping red signals appear in the random coil $^{1}$H chemical shift range between 7.7 and 8.5 ppm, indicating an unstructured N-terminus. In support, these signals are very large and show narrow line widths. Furthermore, no chemical shift changes of signals from residues in the globular domain are observed as it would occur if the N-terminus would interact.

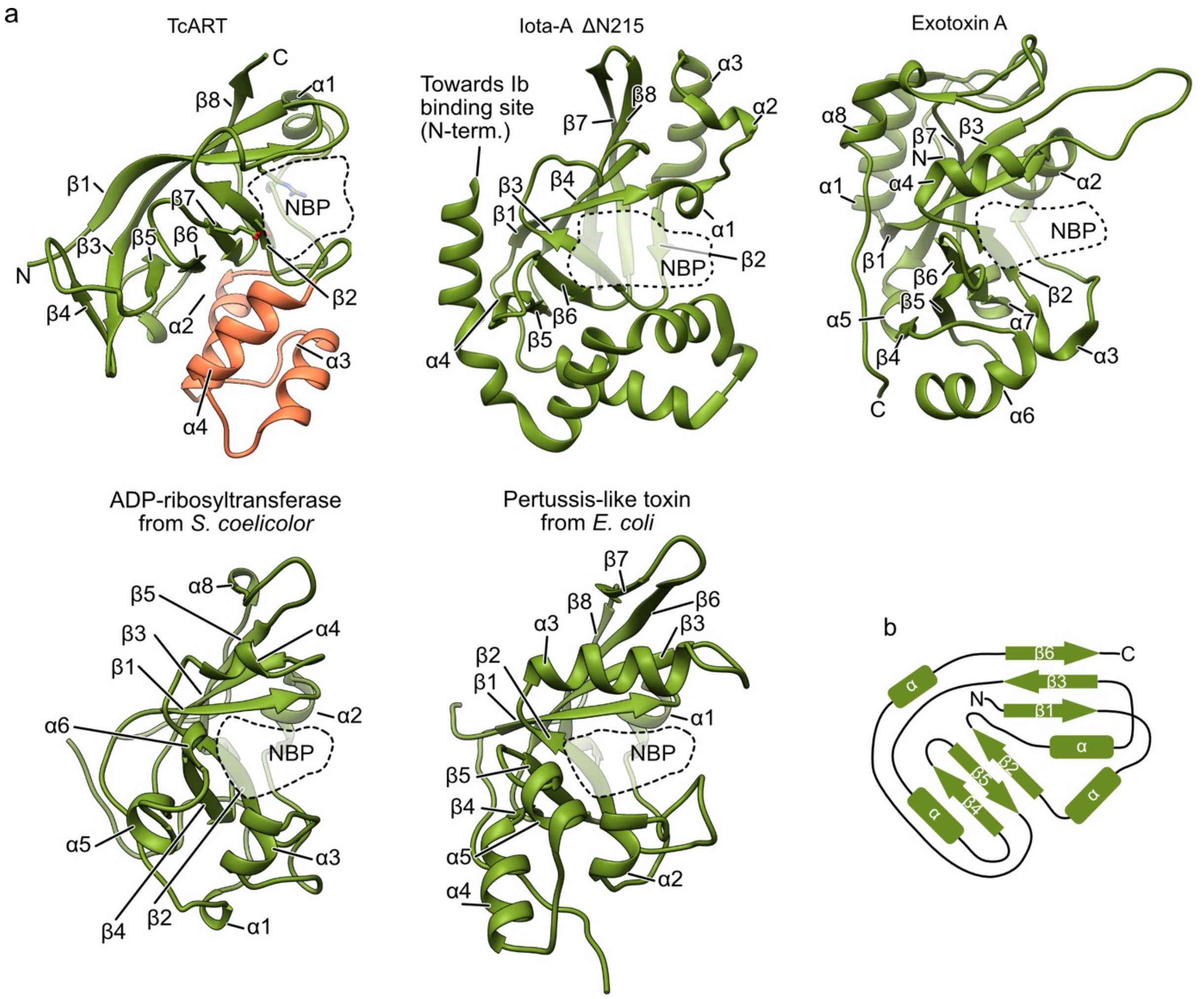

**Fig S3. Comparison of TcART with homologous structures.** (a) Comparison of atomic models of TcART**,** *C. perfringens* Iota toxin (PDB 1GIQ[1]), *P. aeruginosa* Exotoxin A (PDB 3B8H[2]), *S. coelicolor* ADP-ribosyltransferase (PDB 5ZJ5[3]) and *E. coli* pertussis-like toxin (PDB 4Z9D[4]). NBP: nucleotide binding pocket. In the TcART model, R113, S193 and E265 are shown. (b) Average secondary structure of the ART fold according to Aravind et al.[5].

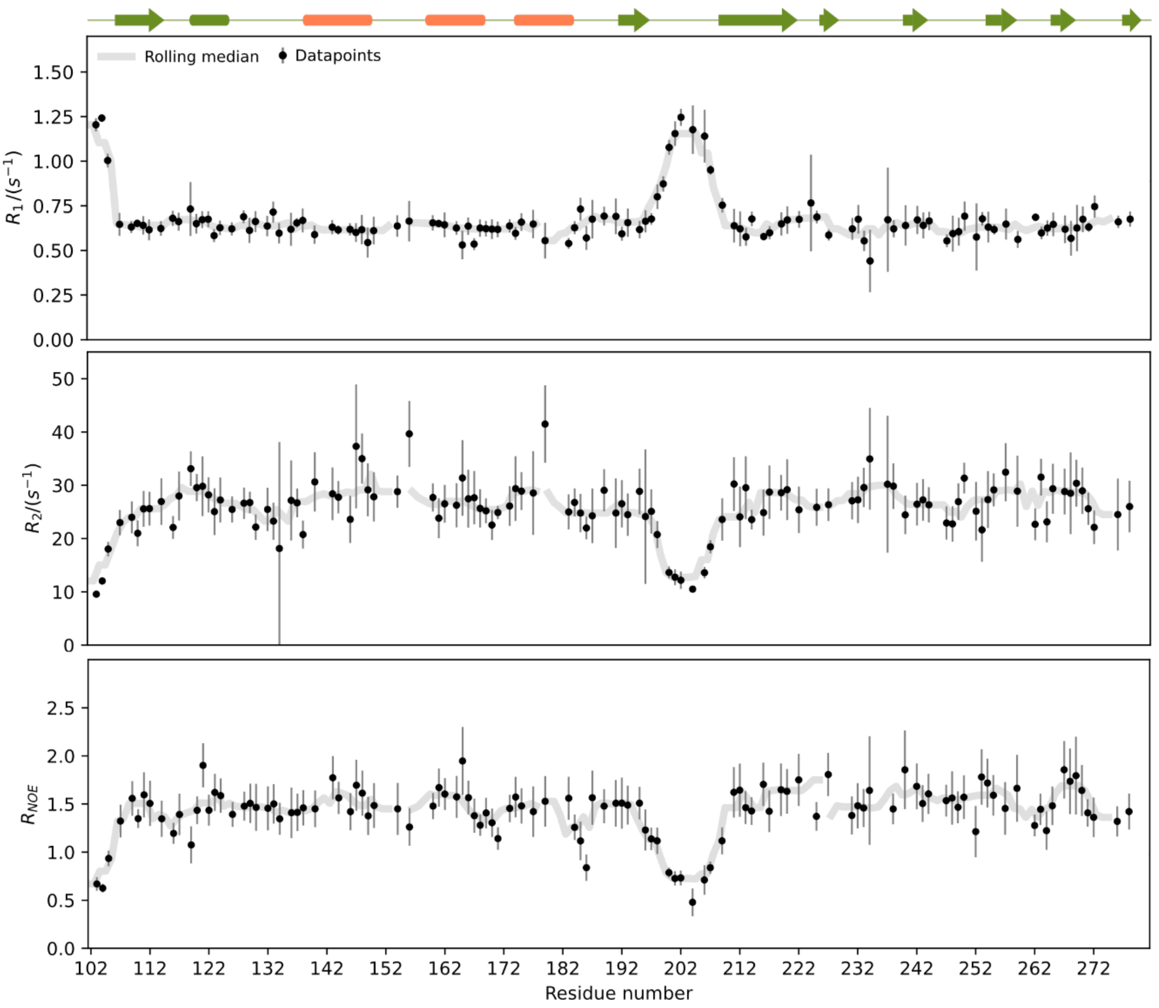

**Fig S4. Longitudinal (top) and transverse (center) $^{15}N$ relaxation rates plotted versus the residue number, together with the $^{1}H$-$^{15}N$-NOE (bottom).** The secondary structure of TcART is given on top. Apart from the values of the N-terminal residues and those in the region 197-208 which indicate high flexibility the protein appears largely as a rigid, globular domain. Error bars indicate twice the standard deviation.

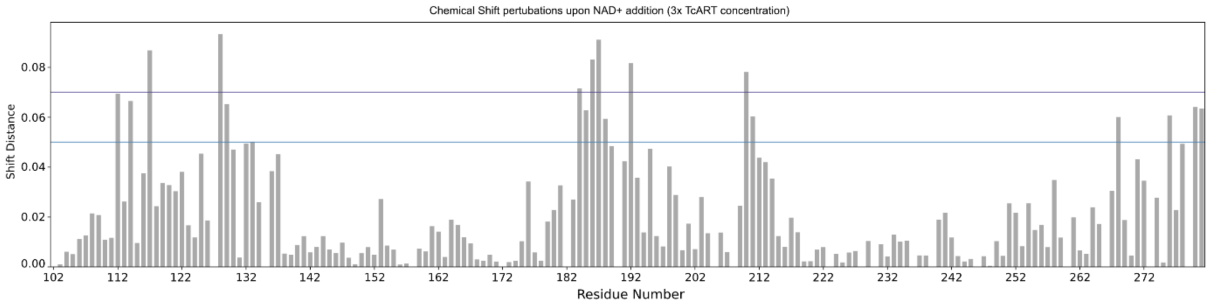

**Fig S5. Chemical shift perturbations per residue within two-dimensional $^{1}H$-$^{15}N$ correlation spectra of TcART upon addition of a three-fold molar excess of $NAD^{+}$.** The values above the blue and lilac lines are the basis for the surface coloring in Fig 1d. Perturbations were quantified as a dimensionless number according to the formula given in the materials and methods part.

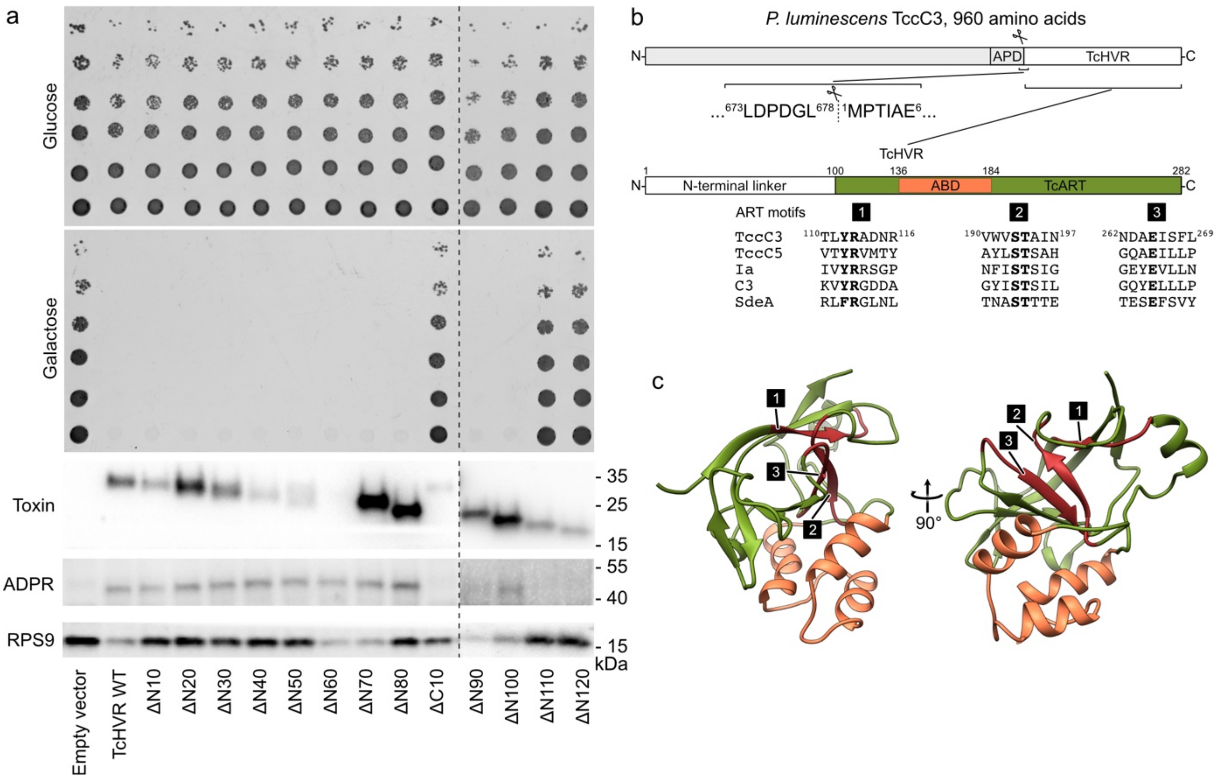

**Fig S6. TcHVR is composed of the N-terminal linker and the C-terminal TcART.** (a) Growth phenotype assay with *S. cerevisiae* expressing TcHVR variants under galactose promoter in the experimental conditions with low (Glucose) or high (Galactose) toxin expression. Analysis of protein expression and actin ADP-ribosylation was performed by western blot of cells grown on galactose-containing media with anti-myc (toxin), anti-ADP-ribose binding reagent (ADPR) and anti-ribosomal protein S9 (RPS9) antibodies. The panel is composed of two drop-test and western blot images as indicated by dashed lines. (b) Schematic representation of the full-length TccC3, TcHVR and TcART. Sequences of 3 conserved ART motifs are provided for *P. luminescence* TccC3 and TccC5, *C. perfringens* Iota toxin, *Bacillus cereus* C3 toxin, *Legionella pneumophila* SdeA. APD – Aspartic protease domain. (c) Positions of ART motifs on the structure of TcART.

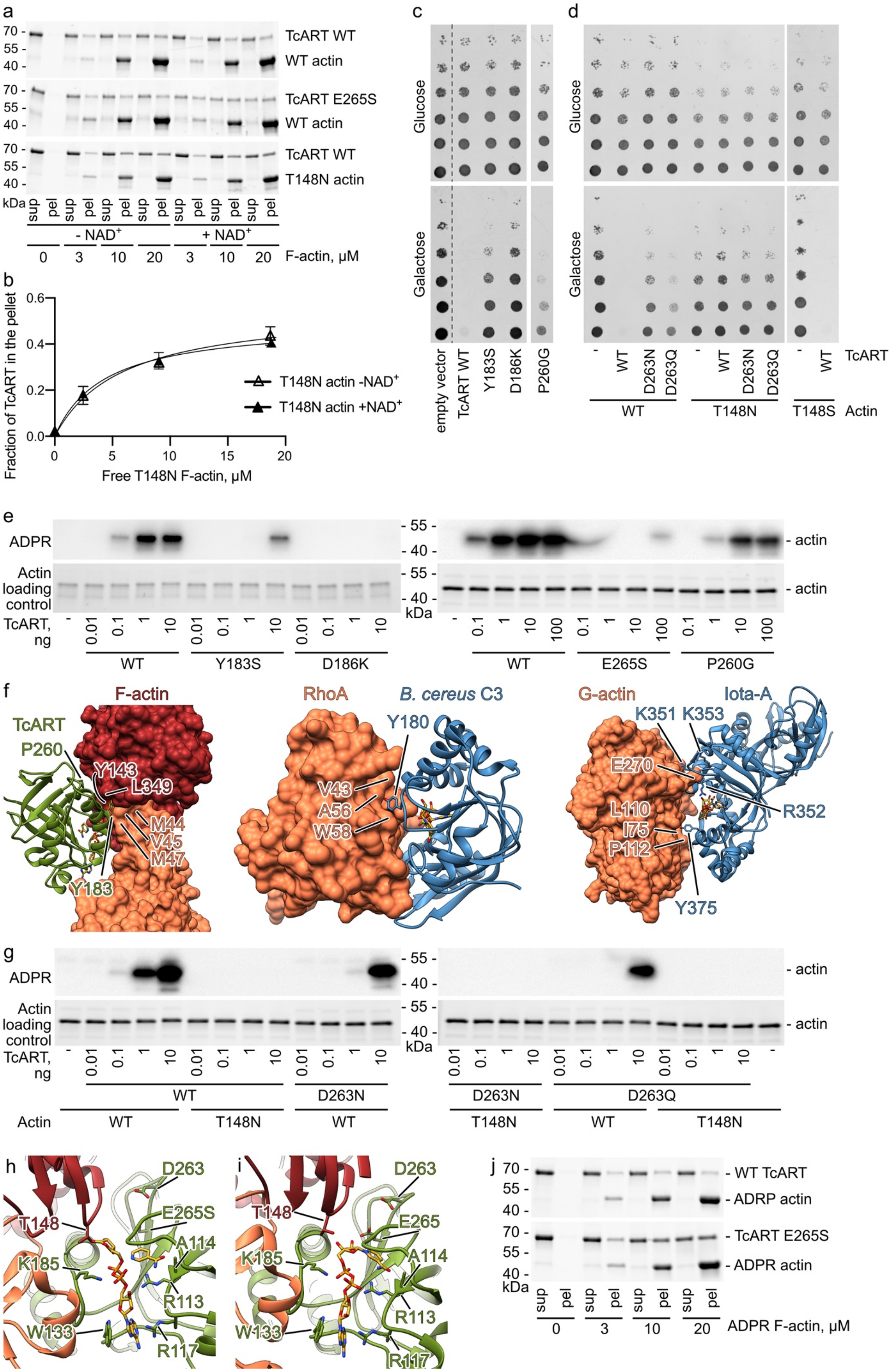

a
TcART WT
WT actin
TcART E265S
WT actin
TcART WT
T148N actin
kDa
sup
pel
- NAD+
+ NAD+
F-actin, μM
b
Fraction of TcART in the pellet
Free T148N F-actin, μM
T148N actin -NAD+
T148N actin +NAD+
c
Glucose
Galactose
empty vector
TcART WT
Y183S
D186K
P260G
d
TcART
WT
D263N
D263Q
Actin
T148N
T148S
e
ADPR
Actin loading control
TcART, ng
actin
E265S
f
TcART
F-actin
P260
Y143
L349
M44
V45
M47
Y183
RhoA
B. cereus C3
Y180
V43
A56
W58
G-actin
Iota-A
K351
K353
E270
R352
L110
I75
P112
Y375
g
h
T148
D263
E265S
A114
K185
R113
W133
R117
i
E265
j
WT TcART
ADRP actin
TcART E265S
ADPR actin
ADPR F-actin, μM

**Fig S7. TcART-F-actin complex.** (a) Co-sedimentation of phalloidin-stabilized F-actin (WT – rabbit muscle α actin, T148N – human cytosolic β actin) and 3 μM TcART analyzed by SDS-PAGE. The upper band corresponds to TcART variants, and the lower band corresponds to F-actin. Representative stain-free gels are shown. (b) The fractions of TcART that co-sedimented with F-actin were quantified by densitometry and plotted against F-actin concentrations. The data are presented as mean values, the error bars correspond to standard deviations of 3 independent experiments. (c) Growth phenotype assay with *S. cerevisiae* expressing WT actin and TcART variants under a strong galactose promoter in the experimental conditions with low (Glucose) or high (Galactose) toxin expression. (d) Growth phenotype assay with *S. cerevisiae* where WT actin was substituted to T148S and T148N variants expressing TcART variants under a strong galactose promoter in the experimental conditions with low (Glucose) or high (Galactose) toxin expression. (e), (g) ADP-ribosylation (ADPR) of 1 μg of phalloidin-stabilized human cytosolic β-F-actin by TcART variants. The reaction mixture was first separated by SDS-PAGE (lower panel), blotted and developed with anti-ADPR reagent in western blot (upper panel). The experiments were performed twice. (f) Comparison of interfaces of the TcART-F-actin, *B. cereus* C3-RhoA (PDB 5BWM[6]) and *C. perfringens* Iota-A-G-actin (PDB 4H03[7]) complexes. (h) Cryo-EM model of the catalytic center of TcART in the post-reaction state. (i) Model of the catalytic center of TcART in the pre-reaction state. The model was obtained by docking of $NAD^+$ to the cryo-EM structure in the post-reaction state. (j) Co-sedimentation of pre-ADP-ribosylated phalloidin-stabilized rabbit muscle F-actin and 3 μM TcART analyzed by SDS-PAGE. The upper band corresponds to TcART variants, and the lower band corresponds to F-actin. Representative stain-free gels are shown. The average values are presented at Fig 3e.

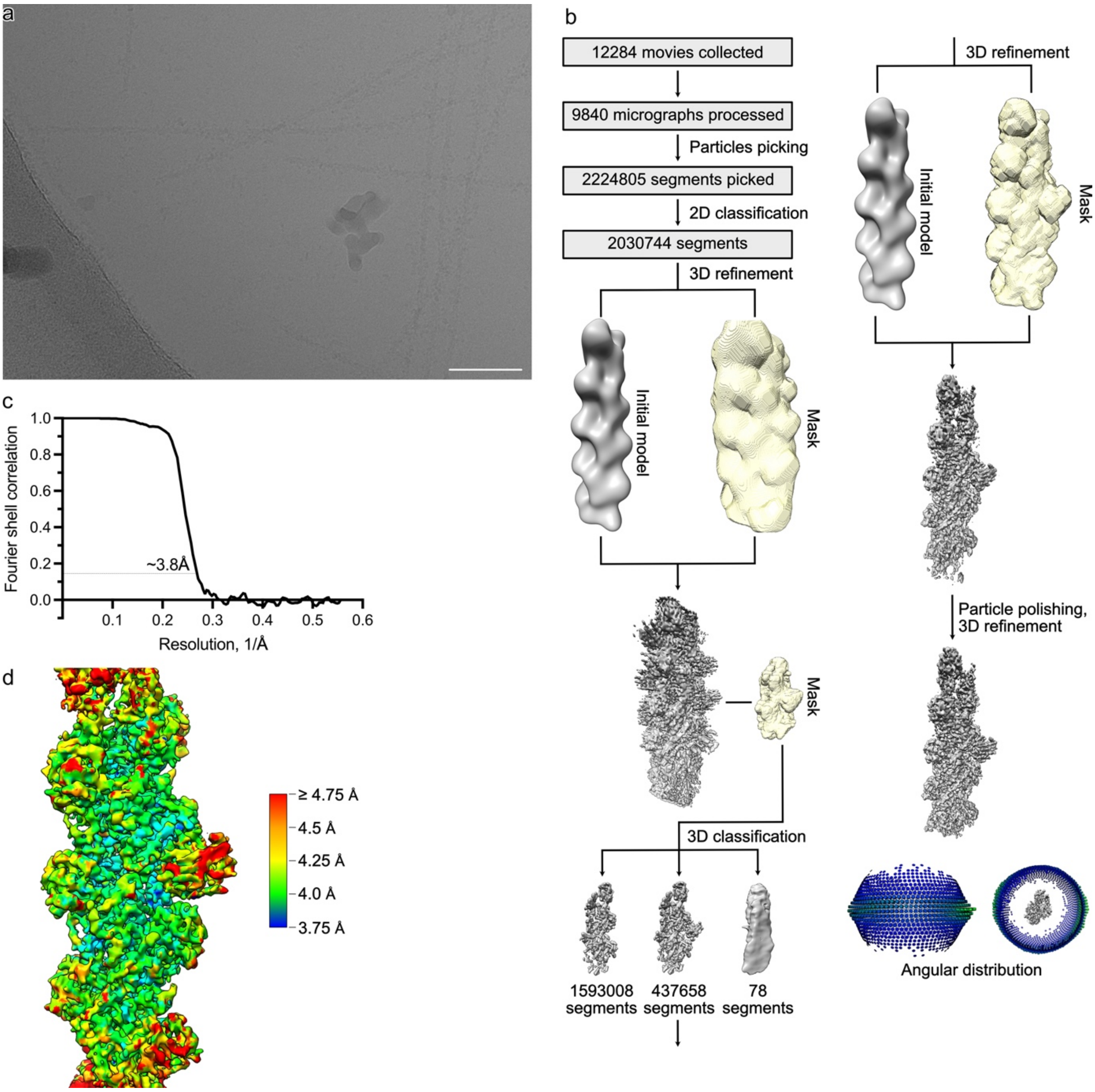

**Fig S8. Processing of the TcART-F-actin complex.** (a) A representative cryo-EM micrograph at 2 µm defocus. Scale bar 50 nm. (b) Processing overview. (c) Fourier shell correlation curve of the final masked map calculated at 120-Å-long central section of the filament. (d) Local resolution gradient of the reconstruction.

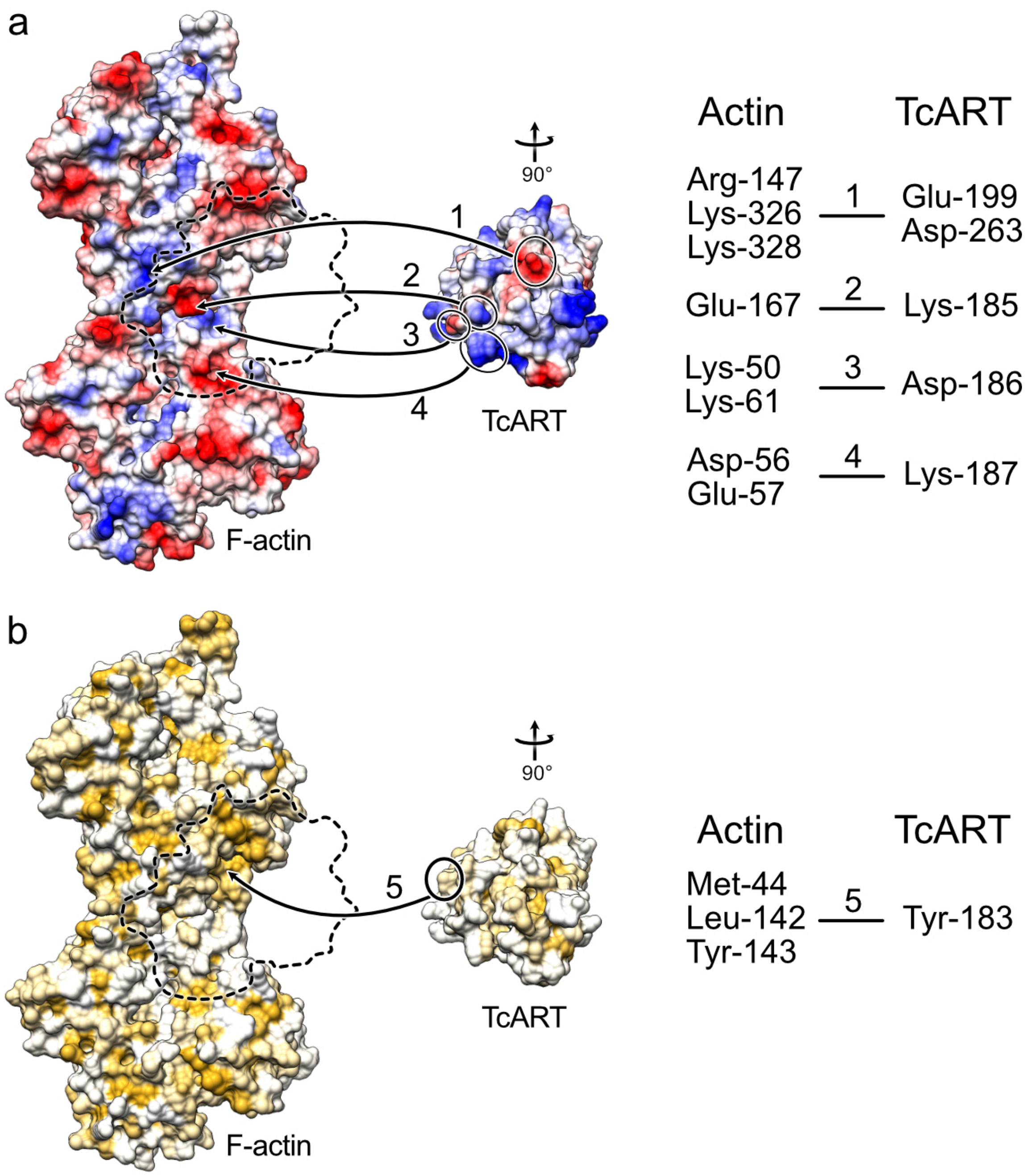

**Fig S9. Interactions in the TcART-F-actin complex.** (a) Electrostatic and (b) hydrophobic interactions in the TcART-F-actin complex.

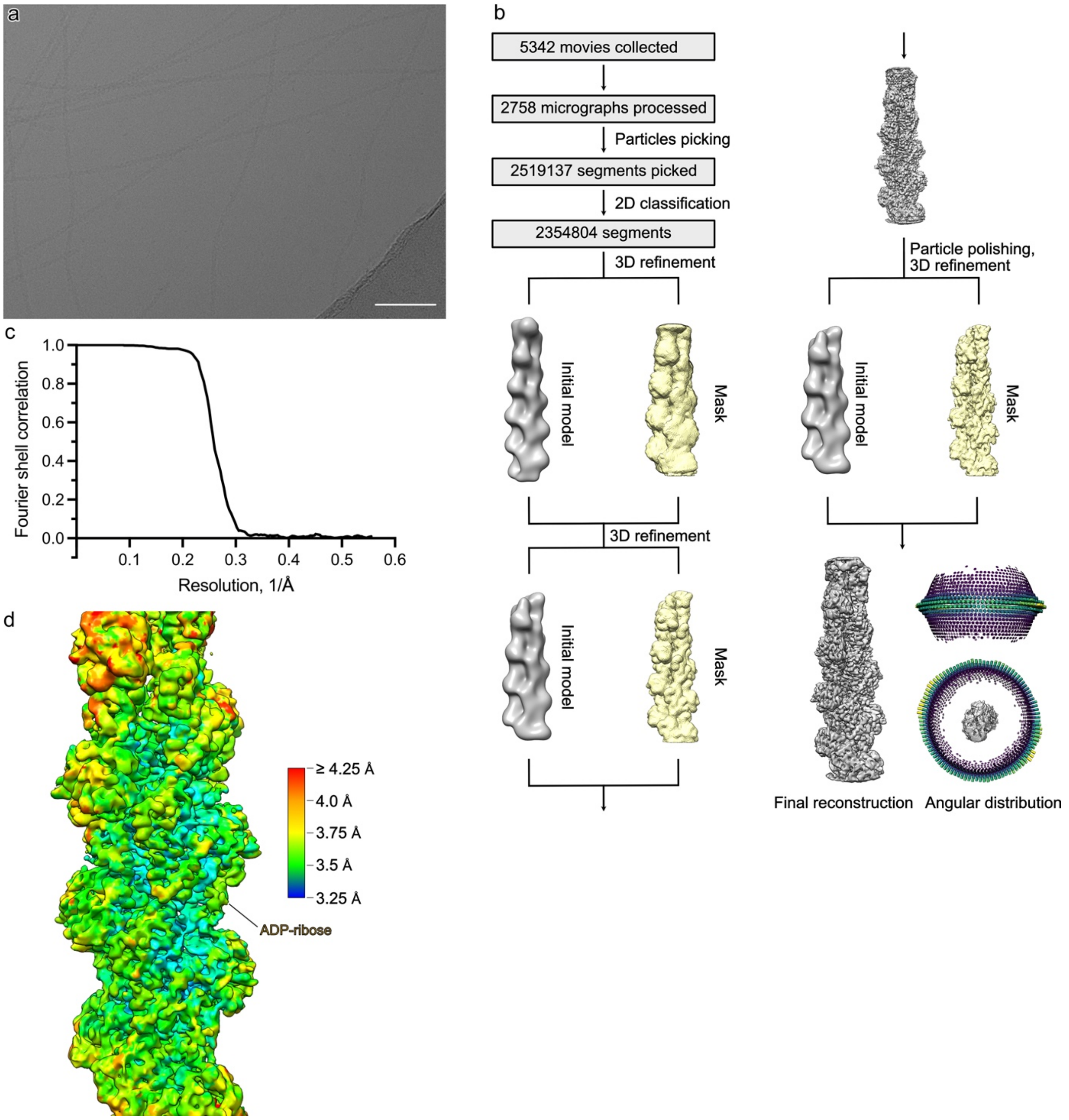

**Fig S10. Processing of the ADP-ribosylated F-actin.** (a) A representative cryo-EM micrograph at 2.5 µm defocus. Scale bar 50 nm. (b) Processing overview. (c) Fourier shell correlation curve of the final masked map calculated at 120-Å-long central section of the filament. (d) Local resolution gradient of the reconstruction.

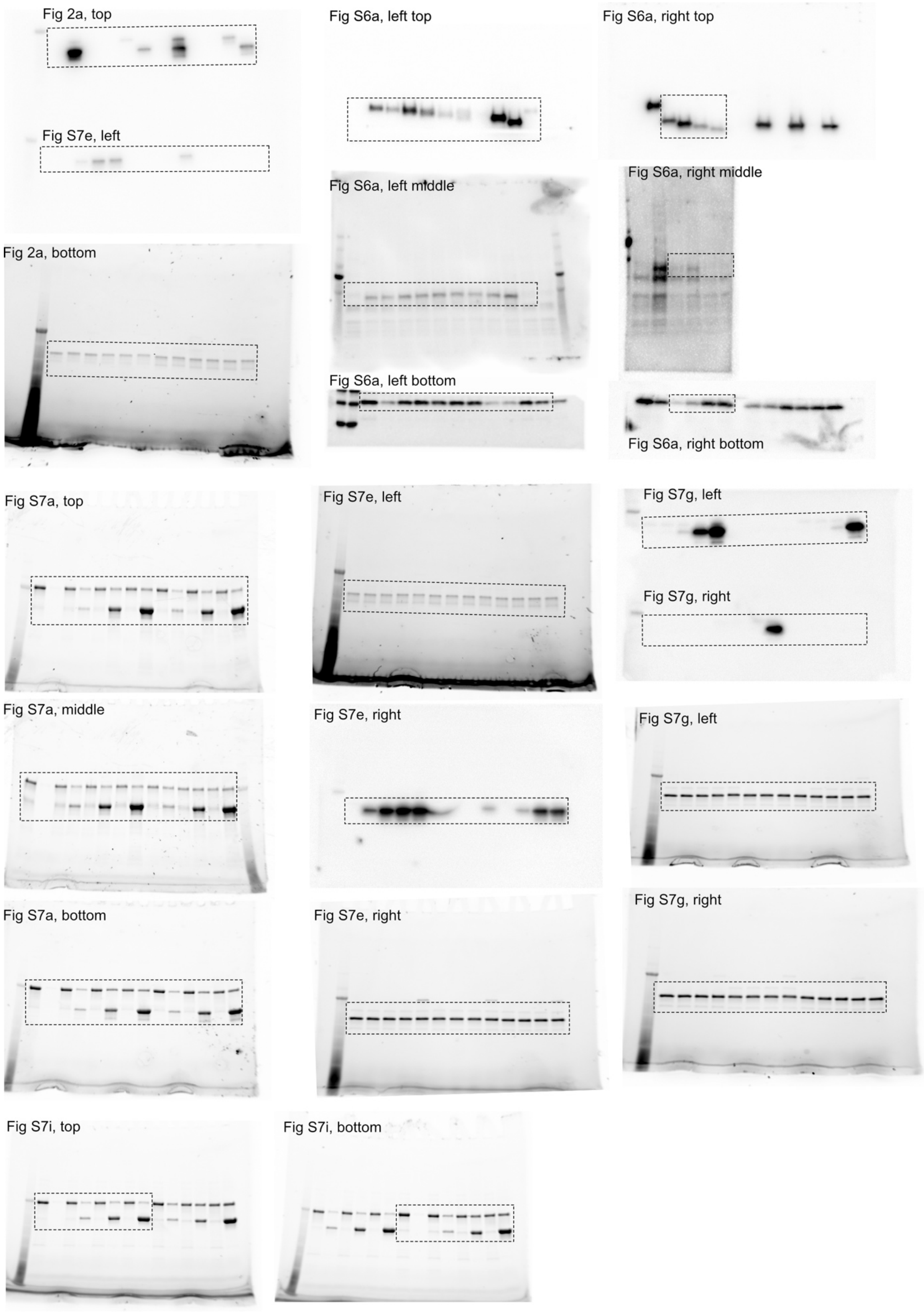

**Fig S11. Uncropped gels and Western blots.**

**Table S1.** Integrals for indicated regions in Figure S1.

| Ref. Area | Estimated Peak amount | Area | absolute Integral | Estimated Peaks | | |
|---|---|---|---|---|---|---|
| 1 | 30 | 1 | 5.50E+12 | 30.00 | | |
| | | 2 | 2.16E+13 | 117.55 | | |
| | | - | exterior Peaks | 2 | Σ | 149.55 |
| | | 3 | 1.62E+12 | 8.84 | | |
| | | 4 | 2.87E+12 | 15.64 | | |
| | | 5 | 1.80E+13 | 98.13 | | |
| | | 6 | 1.19E+12 | 6.49 | | |
| | | 7 | 6.46E+11 | 3.52 | | |
| | | - | exterior Peaks | 12 | Σ | 144.62 |
| 3 | 9 | 1 | 5.50E+12 | 30.55 | | |
| | | 2 | 2.16E+13 | 119.69 | | |
| | | - | exterior Peaks | 2 | Σ | 152.24 |
| | | 3 | 1.62E+12 | 9.00 | | |
| | | 4 | 2.87E+12 | 15.92 | | |
| | | 5 | 1.80E+13 | 99.92 | | |
| | | 6 | 1.19E+12 | 6.61 | | |
| | | 7 | 6.46E+11 | 3.59 | | |
| | | - | exterior Peaks | 12 | Σ | 147.04 |
| 4 | 16 | 1 | 5.50E+12 | 30.70 | | |
| | | 2 | 2.16E+13 | 120.27 | | |
| | | - | exterior Peaks | 2 | Σ | 152.97 |
| | | 3 | 1.62E+12 | 9.04 | | |
| | | 4 | 2.87E+12 | 16.00 | | |
| | | 5 | 1.80E+13 | 100.41 | | |
| | | 6 | 1.19E+12 | 6.65 | | |
| | | 7 | 6.46E+11 | 3.60 | | |
| | | - | exterior Peaks | 12 | Σ | 147.70 |

**Table S2. Cryo-EM data collection, refinement, and validation statistics**

| Project | TcART-F-α-rabbit actin complex | ADPR-F-α-rabbit actin |
|---|---|---|
| Microscope | Titan Krios | |
| Voltage (kV) | 300 | |
| Defocus range (μm) | -0.5 to -2.5 | |
| Camera | Gatan K3 (Superresolution mode) | |
| Pixel size (Å) | 0.9 | |
| Total electron dose (e/Å$^2$) | 84.9 | 82.3 |
| Exposure time (s) | 4 | |
| Frames per movie | 60 | |
| Number of movies | 9840 (12284) | 2758 (5342) |
| **3D Refinement** | | |
| Number of particles | 437658 | 2354804 |
| Final resolution (Å) | 3.8 | 3.5 |
| Helical rise (Å) | 28.1 | 28.2 |
| Helical twist (°) | -167.1 | -166.7 |
| **Atomic model statistics** | | |
| Non-hydrogen atoms | 16102 | 14815 |
| Molprobity score | 2.00 | 1.55 |
| Clashscore | 16.81 | 3.75 |
| EMRinger score | 2.3 | 3.12 |
| Bond RMSD (Å) | 0.01 | 0.011 |
| Angle RMSD (°) | 1.29 | 1.18 |
| Poor rotamers (%) | 0.06 | 0 |
| Favored rotamers (%) | 96.47 | 97.44 |
| Ramachandran favored (%) | 96.47 | 94.26 |
| Ramachandran allowed (%) | 3.43 | 5.74 |
| Ramachandran outliers (%) | 0.1 | 0 |

**Table S3**. **List of primers, strains and plasmids used in this study.**

| Bacterial and yeast strains | Description | Reference |
|---|---|---|
| *E. coli* DH5α | F$^-$ Φ80*lac*ZΔM15 Δ(*lac*ZYA-*arg*F) U169 *rec*A1 *end*A1 *hsd*R17($r_k^-$, $m_k^+$) *pho*A *sup*E44 *thi*-1 *gyr*A96 *rel*A1 λ$^-$ | Invitrogen |
| *E. coli* BL21 DE3 CodonPlus RIPL | F$^-$ *ompT hsdS*($r_B^-$ $m_B^-$) *dcm*$^+$ Tet$^r$ *gal*λ(DE3) *endA* Hte [*argU proL* Cam$^r$] [*argU ileY leuW* Strep/Spec$^r$] | Agilent |
| *S. cerevisiae* MH272-3fα | "Wild-type" strain, *ura3, leu2, his3, trp1, ade2* | [8] |
| *S. cerevisiae* SC483 | *S. cerevisiae* MH272-3fα *act1::*LEU2 + *ACT1*[Ura3] | [9] |
| *S. cerevisiae* SC489 | *S. cerevisiae* MH272-3fα *act1::*LEU2 + *ACT1*[His3] | [9] |
| *S. cerevisiae* Y395 | *S. cerevisiae* MH272-3fα + empty vector [Ade] (2473) | [10] |
| *S. cerevisiae* Y547 | *S. cerevisiae* MH272-3fα + WT TcHVR [Ade] (pB507) | This study |
| *S. cerevisiae* Y617 | *S. cerevisiae* MH272-3fα + TcHVR ΔN10 [Ade] (pB596) | This study |
| *S. cerevisiae* Y618 | *S. cerevisiae* MH272-3fα + TcHVR ΔN20 [Ade] (pB597) | This study |
| *S. cerevisiae* Y619 | *S. cerevisiae* MH272-3fα + TcHVR ΔN30 [Ade] (pB598) | This study |
| *S. cerevisiae* Y620 | *S. cerevisiae* MH272-3fα + TcHVR ΔN40 [Ade] (pB599) | This study |
| *S. cerevisiae* Y621 | *S. cerevisiae* MH272-3fα + TcHVR ΔN50 [Ade] (pB600) | This study |
| *S. cerevisiae* Y622 | *S. cerevisiae* MH272-3fα + TcHVR ΔN60 [Ade] (pB601) | This study |
| *S. cerevisiae* Y623 | *S. cerevisiae* MH272-3fα + TcHVR ΔN70 [Ade] (pB602) | This study |
| *S. cerevisiae* Y624 | *S. cerevisiae* MH272-3fα + TcHVR ΔN80 [Ade] (pB603) | This study |
| *S. cerevisiae* Y638 | *S. cerevisiae* MH272-3fα + TcHVR ΔN90 [Ade] (pB630) | This study |
| *S. cerevisiae* Y639 | *S. cerevisiae* MH272-3fα + TcHVR ΔN100 (ART) [Ade] (pB631) | This study |
| *S. cerevisiae* Y640 | *S. cerevisiae* MH272-3fα + TcHVR ΔN110 [Ade] (pB632) | This study |
| *S. cerevisiae* Y641 | *S. cerevisiae* MH272-3fα + TcHVR ΔN120 [Ade] (pB633) | This study |
| *S. cerevisiae* Y625 | *S. cerevisiae* MH272-3fα + TcHVR ΔC10 [Ade] (pB604) | This study |
| *S. cerevisiae* Y626 | *S. cerevisiae* MH272-3fα + TcHVR ΔC20 [Ade] (pB605) | This study |
| *S. cerevisiae* Y733 | *S. cerevisiae* MH272-3fα + TcART Y183S [Ade] (pB751) | This study |
| *S. cerevisiae* Y735 | *S. cerevisiae* MH272-3fα + TcART D186K [Ade] (pB753) | This study |
| *S. cerevisiae* Y738 | *S. cerevisiae* MH272-3fα *act1::*LEU2 + *ACT1* T148S [His3] (pB748) | This study |
| *S. cerevisiae* Y739 | *S. cerevisiae* MH272-3fα *act1::*LEU2 + *ACT1* T148N [His3] (pB749) | This study |
| *S. cerevisiae* Y740 | *S. cerevisiae* MH272-3fα *act1::*LEU2 + *ACT1* [His3] + empty vector [Ade] (pB87) | This study |
| *S. cerevisiae* Y741 | *S. cerevisiae* MH272-3fα *act1::*LEU2 + *ACT1* [His3] + TcART [Ade] (pB631) | This study |
| *S. cerevisiae* Y742 | *S. cerevisiae* MH272-3fα *act1::*LEU2 + *ACT1* T148S[His3] + empty vector [Ade] (2473) | This study |
| *S. cerevisiae* Y743 | *S. cerevisiae* MH272-3fα *act1::*LEU2 + *ACT1* T148S[His3] + TcART [Ade] (pB631) | This study |
| *S. cerevisiae* Y744 | *S. cerevisiae* MH272-3fα *act1::*LEU2 + *ACT1* T148N [His3] + empty vector [Ade] (2473) | This study |

| | | |
|---|---|---|
| *S. cerevisiae* Y745 | *S. cerevisiae* MH272-3fα *act1::*LEU2 + *ACT1* T148N [His3] + TcART [Ade] (pB631) | This study |
| *S. cerevisiae* Y796 | *S. cerevisiae* MH272-3fα *act1::*LEU2 + *ACT1*[His3] + TcART P260G [Ade] (pB849) | This study |
| *S. cerevisiae* Y797 | *S. cerevisiae* MH272-3fα *act1::*LEU2 + *ACT1*[His3] + TcART D263N [Ade] (pB850) | This study |
| *S. cerevisiae* Y798 | *S. cerevisiae* MH272-3fα *act1::*LEU2 + *ACT1*[His3] + TcART D263Q [Ade] (pB851) | This study |
| *S. cerevisiae* Y800 | *S. cerevisiae* MH272-3fα *act1::*LEU2 + *ACT1* T148N [His3] + TcART D263N [Ade] (pB850) | This study |
| *S. cerevisiae* Y801 | *S. cerevisiae* MH272-3fα *act1::*LEU2 + *ACT1* T148N [His3] + TcART D263Q [Ade] (pB851) | This study |
| **Plasmids for experiments in *S. cerevisiae*** | | |
| 2473 YEpGal555 | *E. coli*/*S. cerevisiae* shuttle vector [ADE2] with Gal1 promoter | [9] |
| pB507 TcHVR WT | TcHVR WT was amplified from p579 using oligonucleotides tatactcgagttaatgccaacaattgcagaacg and tatatagctagctctcttatgaggttttacatttttaagc. The PCR product was digested with XhoI and NheI and ligated into digested 2473 vector. | This study |
| pB596 TcHVR ΔN10 | TcHVR with the N-terminal deletion was amplified from pB501 using oligonucleotides tatactcgagctaaaaaaaaataaagtaacagactcagcgcc and tatatagctagctctcttatgaggttttacatttttaagc. The PCR product was digested with XhoI and NheI and ligated into digested 2473 vector. | This study |
| pB597 TcHVR ΔN20 | TcHVR with the N-terminal deletion was amplified from pB501 using oligonucleotides tatactcgagccttcgccagcaaatgcc and tatatagctagctctcttatgaggttttacatttttaagc. The PCR product was digested with XhoI and NheI and ligated into digested 2473 vector. | This study |
| pB598 TcHVR ΔN30 | TcHVR with the N-terminal deletion was amplified from pB501 using oligonucleotides tatactcgagataaacatccgccccgcctgtag and tatatagctagctctcttatgaggttttacatttttaagc. The PCR product was digested with XhoI and NheI and ligated into digested 2473 vector. | This study |
| pB599 TcHVR ΔN40 | TcHVR with the N-terminal deletion was amplified from pB501 using oligonucleotides tatactcgagcctagcttaccgaaagcatcaacg and tatatagctagctctcttatgaggttttacatttttaagc. The PCR product was digested with XhoI and NheI and ligated into digested 2473 vector. | This study |
| pB600 TcHVR ΔN50 | TcHVR with the N-terminal deletion was amplified from pB501 using oligonucleotides tatactcgagcaaccaaccacacaccctatcg and tatatagctagctctcttatgaggttttacatttttaagc. The PCR product was digested with XhoI and NheI and ligated into digested 2473 vector. | This study |
| pB601 TcHVR ΔN60 | TcHVR with the N-terminal deletion was amplified from pB501 using oligonucleotides tatactcgagaacataaaaccaacgacgtctggg and tatatagctagctctcttatgaggttttacatttttaagc. The PCR product was digested with XhoI and NheI and ligated into digested 2473 vector. | This study |
| pB602 TcHVR ΔN70 | TcHVR with the N-terminal deletion was amplified from pB501 using oligonucleotides tatactcgagattgttgctccattgagtccagtag and | This study |

| | | |
|---|---|---|
| | tatatagctagctctcttatgaggttttacatttttaagc. The PCR product was digested with XhoI and NheI and ligated into digested 2473 vector. | |
| pB603 TcHVR ΔN80 | TcHVR with the N-terminal deletion was amplified from pB501 using oligonucleotides tatactcgagaaatctacttctgaaatctctctgccagaaag and tatatagctagctctcttatgaggttttacatttttaagc. The PCR product was digested with XhoI and NheI and ligated into digested 2473 vector. | This study |
| pB604 TcHVR ΔC10 | TcHVR with the C-terminal deletion was amplified from pB501 using oligonucleotides tatactcgagttaatgccaacaattgcagaacg and tatatagctagcaattgttgtcagaaatgaaatttctgcatcatttaccgg. The PCR product was digested with XhoI and NheI and ligated into digested 2473 vector. | This study |
| pB605 TcHVR ΔC20 | TcHVR with the C-terminal deletion was amplified from pB501 using oligonucleotides tatactcgagttaatgccaacaattgcagaacg and tatatagctagcatttaccggtccatgattaagtgcaatg. The PCR product was digested with XhoI and NheI and ligated into digested 2473 vector. | This study |
| pB630 TcHVR ΔN90 | TcHVR with the N-terminal deletion was amplified from pB501 using oligonucleotides tatactcgagagcgctcaaagcagttcttcaagc and tatatagctagctctcttatgaggttttacatttttaagc. The PCR product was digested with XhoI and NheI and ligated into digested 2473 vector. | This study |
| pB631 TcHVR ΔN100 (TcART) | TcHVR with the N-terminal deletion was amplified from pB501 using oligonucleotides tatactcgagtcgacaaatctacagaaaaaatcatttac and tatatagctagctctcttatgaggttttacatttttaagc. The PCR product was digested with XhoI and NheI and ligated into digested 2473 vector. | This study |
| pB632 TcHVR ΔN110 | TcHVR with the N-terminal deletion was amplified from pB501 using oligonucleotides tatactcgagttatatagagcagataacagatcc and tatatagctagctctcttatgaggttttacatttttaagc. The PCR product was digested with XhoI and NheI and ligated into digested 2473 vector. | This study |
| pB633 TcHVR ΔN120 | TcHVR with the N-terminal deletion was amplified from pB501 using oligonucleotides tatactcgaggaaatgcaaagtaaattccctgaagg and tatatagctagctctcttatgaggttttacatttttaagc. The PCR product was digested with XhoI and NheI and ligated into digested 2473 vector. | This study |
| pB748 pRS313 Actin T148S | The T148S mutation was generated by two-step overlap PCR using oligonucleotides ctcttccggtagatctactggtattg, caggaaacagctatgacc, caataccagtagatctaccggaagag and ttcgtgataagtgatagtg. The PCR product was digested with ClaI and SalI and was used to exchange the WT ACT1 gene in 2475 p1387 pRS313 Actin [9]. | This study |
| pB749 pRS313 Actin T148N | The T148N mutation was generated by two-step overlap PCR using oligonucleotides gtactcttccggtagaaacactggtattgttttgg, caggaaacagctatgacc, ccaaaacaataccagtgtttctaccggaagagtac and ttcgtgataagtgatagtg. The PCR product was digested with ClaI and SalI and was used to exchange the WT ACT1 gene in 2475 p1387 pRS313 Actin [9]. | This study |
| pB751 TcART Y183S | The Y183S mutation was generated by two-step overlap PCR using oligonucleotides tatactcgagtcgacaaatctacagaaaaaatcatttac, tatatagctagctctcttatgaggttttacatttttaagc, caaattacataaaatctaccaaggac and | This study |

| | | |
|---|---|---|
| | gtccttggtagattttatgtaatttg. The PCR product was digested with XhoI and NheI and ligated into digested 2473 vector. | |
| pB753 TcART D186K | The D186K mutation was generated by two-step overlap PCR using oligonucleotides tatactcgagtcgacaaatctacagaaaaaatcatttac, tatatagctagctctcttatgaggttttacatttttaagc, cataaaatataccaagaaaaaatctacagtatg and catactgtagattttttcttggtatattttatg. The PCR product was digested with XhoI and NheI and ligated into digested 2473 vector. | This study |
| pB849 TcART P260Q | TcART with the P260Q mutation was amplified from pB631 using oligonucleotides tatactcgagtcgacaaatctacagaaaaaatcatttac and atcttagctagctctcttatgaggttttacatttttaagcggaattgttgtcagaaatgaaatttctgcatcatttactcctccatgattaag. The PCR product was digested with XhoI and NheI and ligated into digested 2473 vector. | This study |
| pB850 TcART D263N | TcART with the D263N mutation was amplified from pB631 using oligonucleotides tatactcgagtcgacaaatctacagaaaaaatcatttac and atcttagctagctctcttatgaggttttacatttttaagcggaattgttgtcagaaatgaaatttctgcattatttaccgg. The PCR product was digested with XhoI and NheI and ligated into digested 2473 vector. | This study |
| pB851 TcART D263Q | TcART with the D263Q mutation was amplified from pB631 using oligonucleotides tatactcgagtcgacaaatctacagaaaaaatcatttac and atcttagctagctctcttatgaggttttacatttttaagcggaattgttgtcagaaatgaaatttctgcttgatttaccgg. The PCR product was digested with XhoI and NheI and ligated into digested 2473 vector. | This study |
| **Plasmids for protein expression in *E. coli*** | | |
| 1330 pET28 Iota-A | *Clostridium perfringens* Iota-A with N-terminal His-tag in pET28 vector | [9] |
| 579 pET19 TcHVR WT | TcHVR WT with C-terminal His-tag in pET19 vector | [11] |
| 613 TcdB2-TccC3 | Fusion construct of TcdB2–TccC3 | [12] |
| pB137 pET28 MBP | Maltose-binding protein with N-terminal His-tag in pET28 vector | [10] |
| pB656 pET28 MBP-TcHVR ΔN100 (TcART) | TcHVR with the N-terminal deletion was amplified from p579 using oligonucleotides tatagagctctggttcgacaaatctacagaaaaaatcatttac and tataaagcttttatctcttatgaggttttacatttttaagc. The PCR product was digested with SacI and HindIII and ligated into digested pB137 vector. | This study |
| pB685 pET28 MBP-TcART E265S | The E265S mutation was generated by two-step overlap PCR using oligonucleotides tatagagctctggttcgacaaatctacagaaaaaatcatttac, tataaagcttttatctcttatgaggttttacatttttaagc, gtaaatgatgcatcaatttcatttc and gaaatgaaattgatgcatcatttac. The PCR product was digested with SacI and HindIII and ligated into digested pB137 vector. | This study |
| pB751 pET28 MBP-TcART Y183S | The Y183S mutation was generated by two-step overlap PCR using oligonucleotides tatagagctctggttcgacaaatctacagaaaaaatcatttac, tataaagcttttatctcttatgaggttttacatttttaagc, caaattacataaaatctaccaaggac and gtccttggtagattttatgtaatttg. The PCR product was digested with SacI and HindIII and ligated into digested pB137 vector. | This study |
| pB753 pET28 MBP-TcART D186K | The D186K mutation was generated by two-step overlap PCR using oligonucleotides tatagagctctggttcgacaaatctacagaaaaaatcatttac, | This study |

| | | |
|---|---|---|
| | tataaagcttttatctcttatgaggttttacatttttaagc, cataaaatataccaagaaaaaatctacagtatg and catactgtagattttttcttggtatattttatg. The PCR product was digested with SacI and HindIII and ligated into digested pB137 vector. | |
| pB846 pET28 MBP-TcART P260Q | TcART with the P260Q mutation was amplified from pB656 using oligonucleotides tatagagctctggttcgacaaatctacagaaaaaatcatttac and tataaagcttttatctcttatgaggttttacatttttaagcggaattgttgtcagaaatgaaatttctgcatcattta cacctccatgattaag. The PCR product was digested with SacI and HindIII and ligated into digested pB137 vector. | This study |
| pB847 pET28 MBP-TcART D263N | TcART with the D263N mutation was amplified from pB656 using oligonucleotides tatagagctctggttcgacaaatctacagaaaaaatcatttac and tataaagcttttatctcttatgaggttttacatttttaagcggaattgttgtcagaaatgaaatttctgcattattta ccggtc. The PCR product was digested with SacI and HindIII and ligated into digested pB137 vector. | This study |
| pB848 pET28 MBP-TcART D263Q | TcART with the D263Q mutation was amplified from pB656 using oligonucleotides tatagagctctggttcgacaaatctacagaaaaaatcatttac and tataaagcttttatctcttatgaggttttacatttttaagcggaattgttgtcagaaatgaaatttctgcttgattta ccggtc. The PCR product was digested with SacI and HindIII and ligated into digested pB137 vector. | This study |
| 2013 pET19b cHis-3C TcHVR ΔN100 (TcART) | TcHVR with the N-terminal deletion was amplified from p579 and cloned into pET19b vector. | This study |
| **Plasmids for protein expression in insect cells** | | |
| 2336 pFL ACTB C272A | Human ActB with C272A mutation followed by the cleavable linker and thymosin β4 | [13] |
| 2723 pFL ACTB T148N C272A | T148N mutations was introduced by the QuikChange method using oligonucleotides ctgtacgcctctggccgtaacactggcatcgtgatggactcc and ggagtccatcacgatgccagtgttacggccagaggcgtacag and 2336 as a matrix | This study |

**Table S4.** Parameters of NMR data acquisition and processing.

| | Experiment | MHz | (F3 x )F2 x F1* | scans | complex datapoints | d1 in s | expt time | aq time in ms | SW in ppm | mixing time | processing datapoints | processing window functions |
|---|---|---|---|---|---|---|---|---|---|---|---|---|
| 3D | CCH TOCSY | 600 | H x C x C | 16 | 1024 x 128 x 128 | 1 | 3d 8h | 51.2 x 5.2 x 5.2 | 16.7 x 80.8 x 80.8 | 10 ms | 2048 x 512 x 256 | qsine 2.5 x qsine 2 x qsine 2 |
| 2D | CH $D_2O$ | 600 | H x C | 16 | 1024 x 256 | 1.3 | 1h 35min | 51.2 x 10.5 | 16.7 x 80.8 | - | 4096 x 2048 | qsine 2 x qsine 2 |
| 3D | CHH NOESY aliphatic | 750 | H x C x H | 8 | 1024 x 164 x 300 | 1.3 | 6d 14h | 41 x 5.6 x 15.0 | 16.7 x 78 x 13.3 | 80 ms | 1024 x 512 x 512 | qsine 2 x qsine 2 x qsine 2 |
| 3D | CHH NOESY aliphatic NAD+ | 600 | H x C x H | 16 | 1024 x 128 x 73 | 1.3 | 2d 12h | 34.1 x 5.5 x 12.1 | 25 x 77 x 15.1 | 80 ms | 2048 x 512 x 512 | qsine 2 x qsine 2 x qsine 2 |
| 3D | CHH NOESY aliphatic $D_2O$ | 600 | H x C x H | 8 | 1024 x 128 x 256 | 1.3 | 3d 18h | 51.2 x 5.2 x 15.4 | 16.7 x 80.8 x 13.9 | 40 ms | 2048 x 512 x 256 | qsine 3 x qsine 2 x qsine 2 |
| 3D | CHH NOESY aromatic | 750 | H x C x H | 8 | 1024 x 150 x 320 | 1.3 | 6d 10h | 41 x 5.1 x 16 | 16.7 x 78 x 13.3 | 80 ms | 2048 x 512 x 1024 | qsine 3 x qsine 2 x qsine 3 |
| 2D | HH NOESY $D_2O$ Ndec | 600 | H x H | 80 | 2048 x 1024 | 1.3 | 1d 10h | 102.4 x 6.1 | 16.7 x 13.9 | 40 ms | 4096 x 4096 | gm (-40.0 \| 0.1) x qsine 3 |
| 2D | HN $D_2O$ | 600 | H x N | 32 | 1024 x 256 | 0.1 | 25 min | 51.2 x 42.5 | 16.7 x 49.5 | - | 4096 x 1024 | qsine 3 x qsine 2 |
| 2D | HN prot | 600 | H x N | 4 | 2048 x 512 | 1.3 | 50 min | 102.4 x 6.1 | 16.7 x 49.5 | - | 4096 x 1024 | qsine 3 x qsine 3 |
| 2D | HN trosy | 600 | H x N | 32 | 1024 x 1024 | 1.3 | 13h 16min | 51.2 x 17 | 16.7 x 49.5 | - | 4096 x 2048 | qsine 2.5 x qsine 2 |
| 3D | HNCA prot | 600 | H x N x C | 32 | 1024 x 128 x 100 | 1.3 | 3d 10h | 51.2 x 21.2 x 7.1 | 16.7 x 49.5 x 46.7 | - | 4096 x 256 x 256 | qsine 3 x qsine 2 x qsine 2 |
| 3D | HNCACB | 600 | H x N x C | 16 | 1024 x 80 x 140 | 1.3 | 3d 7h | 51.2 x 16 x 7 | 16.7 x 41.1 x 66.3 | - | 1024 x 512 x 512 | qsine 2.5 x qsine 2 x qsine 2 |
| 3D | HNCO | 600 | H x N x C | 8 | 1024 x 80 x 128 | 1.3 | 1d 9h | 51.2 x 16 x 25.6 | 16.7 x 41.1 x 16.6 | - | 1024 x 512 x 512 | qsine 2.5 x qsine 2 x qsine 2 |
| 3D | HNCOCACB | 600 | H x N x C | 16 | 1024 x 80 x 140 | 1.3 | 3d 8h | 51.2 x 16 x 7 | 16.7 x 41.1 x 66.3 | - | 1024 x 512 x 512 | qsine 2.5 x qsine 2 x qsine 2 |
| 3D | NHH NOESY 80ms mixing | 750 | H x N x H | 8 | 1024 x 128 x 300 | 1.3 | 5d 4h | 41 x 20.5 x 15 | 16.7 x 41.1 x 13.3 | 80 ms | 2048 x 256 x 512 | qsine 3 x qsine 2 x qsine 3 |
| 2D | NH + 0 mM NAD+ (= HN prot) | 600 | H x N | 4 | 2048 x 512 | 1.3 | 50 min | 102.4 x 6.1 | 16.7 x 49.5 | - | 4096 x 2048 | qsine 3 x qsine 3 |
| 2D | NH + 1 mM NAD+ | 600 | H x N | 4 | 1024 x 256 | 1.3 | 24 min | 51.2 x 42.5 | 16.7 x 49.5 | - | 4096 x 2048 | qsine 3 x qsine 3 |
| pseudo 3D | T1 relaxation | 600 | H x N x t1 | 32 | 1024 x 256 x 8 | 1.3 | 2d 5h | 51.2 x 42.5 x 1500 | 16.7 x 49.5 x 0 | - | 4096 x 1024 | qsine 2 x qsine 2 |
| pseudo 3D | T2 relaxation | 600 | H x N x t1 | 32 | 1024 x 256 x 8 | 1.3 | 1d 6h | 51.2 x 42.5 x 220 | 16.7 x 49.5 x 0 | - | 4096 x 1024 | qsine 2 x qsine 2 |
| pseudo 3D | hetero-NOE | 600 | H x N x NOE | 96 | 1024 x 256 x 2 | 1.3 | 1d 23h | 51.2 x 42.5 x 220 | 16.7 x 49.5 x 0 | - | 4096 x 1024 | qsine 2 x qsine 2 |
| 2D | NH of cocoon without toxin inside | 900 | H x N | 32 | 2048 x 1200 | 0.98 | | 28.7 x 48.0 | 39.7 x 137 | - | 4096 x 4096 | gm (-20 \| 0.1) x qsine 3 |
| | | | | | | | | | | | 4096 x 4096 | gm (-30 \| 0.04) x qsine 2 |
| 2D | NH of cocoon with toxin inside | 900 | H x N | 32 | 2048 x 1200 | 1 | | 28.7 x 48.0 | 39.7 x 137 | - | 4096 x 4096 | gm (-20 \| 0.1) x qsine 3 |
| | | | | | | | | | | | 4096 x 4096 | gm (-30 \| 0.04) x qsine 2 |

*highest dimension is always the direct one

**Table S5**. Summary of structural statistics for the NMR ensemble.

| | |
|---|---|
| Completeness of resonance assignment | |
| Backbone[a] | 91% |
| Sidechain non-H | 74% |
| Sidechain H | 89% |
| Aromatic | 89% |
| Conformationally restricting restraints (consensus ensembles) | |
| Distance restraints | |
| Total | 3315 |
| Intraresidue (i=j) | 0 |
| Sequential (\|i-1\|=1) | 834 |
| Medium range (1<\|i-j\|<5) | 460 |
| Long range (\|i-j\|>=5) | 1297 |
| Ambiguous | 724 |
| Dihedral angle restr. | 306 |
| Hydrogen-bond restr. | 37 |
| Residual restraint violations (average of 2 consensus runs) | |
| Average no. of distance viol. per structure | |
| 0.1 - 0.3 Å | 209 |
| 0.3 - 0.5 Å | 165 |
| > 0.5 Å | 270 |
| Average no. of dihedral viol. per structure | |
| > 5° | 3.6 |
| Model quality[b] | |
| RMSD backbone atoms (Å)[c] | 0.5 |
| RMSD heavy atoms (Å)[c] | 0.8 |
| RMSD bond lenghts (Å) | 0.005 |
| RMSD bond angles (°) | 0.7 |
| Molprobity Ramachandran Statistics[b,c] | |
| Most favored regions (%) | 97.2 |
| Allowed regions (%) | 2.7 |
| Disallowed regions (%) | 0.1 |
| Global quality scores (raw/Z scores)[b] | |
| Verify3D | (0.25/-3.37) |
| Prosall | (0.60/-0.21) |
| Procheck (ϕ-ψ)[c] | (-0.42/-1.34) |
| Procheck (all)[c] | (-0.35/-2.07) |
| MolProbity clash score | (24.08/-2.61) |
| Model contents | |
| Ordered residue ranges[b] | 103-200, 207-231, 237-258, 260-280 |
| Total no. of residues | 180 |
| BMRB accession number | 34717 |
| PDB ID Code | 7ZBQ |

[a]CO resonances were only assigned for the deuterated protein

[b]calculated using the PSVS webserver (https://montelionelab.chem.rpi.edu/PSVS/PSVS/)

[c]for all ordered residues from PSVS based on dihedral order parameter S(ϕ)+S(ψ)>=1.8
